# Two-photon characterisation of long-Stokes-shift dye ATTO 490LS for single-laser multicolour imaging

**DOI:** 10.1101/2025.11.21.689649

**Authors:** King Yee Cheung, Yue Wu, Shu Ying Lee, Xianyuan Zhang, Masahiro Fukuda, Danesha Devini Suresh, Adam Claridge-Chang

## Abstract

Long-Stokes-shift fluorophores enable high sensitivity and multiplexed imaging with single-wavelength excitation. Under single-photon illumination ATTO 490LS exhibits a 165-nm Stokes shift, but its two-photon properties remain uncharacterised. Emission and excitation spectral analyses of ATTO 490LS in ex vivo *Drosophila* melanogaster brains identified two-photon excitation sensitivity at 940 nm, with peak emission at 640 nm. We demonstrate successful duplexed imaging of ATTO 490LS alongside Alexa Fluor 488 using a single 920-nm fibre laser and dual photomultiplier tubes, enabling distinct measurement of red and green fluorescence signals. These findings establish ATTO 490LS as suitable for multicolour two-photon microscopy with single-laser systems.

## Description

A Stokes shift refers to the difference in wavelength between maximum absorption and maximum emission of a fluorophore (Stokes, 1852). A long Stokes shift describes when a fluorescent molecule is excitable by a relatively short wavelength, but emits at a much longer wavelength due to a loss of energy following excitation (Lakowicz, 2001; Valeur, 2001). This can be valuable in fluorescence microscopy, as it reduces overlap between excitation and emission spectra, thereby minimising noise from scattered excitation light and improving detection sensitivity (Sednev et al., 2015; Liu et al., 2024).

Long-Stokes-shift fluorophores can also be suitable for multicolour imaging on microscope systems that are limited to a single laser. A long-Stokes-shift dye can be combined with another fluorophore with the same or a similar excitation profile, but give a completely different emission spectrum, thus making the two dyes distinguishable during image collection (Likhotkin et al., 2023). ATTO 490LS (ATTO-TEC GmbH) is an example of a red-fluorescing long-Stokes-shift dye, with a reported single-photon excitation wavelength of 496 nm and emission wavelength of 661 nm (Stokes shift of 165 nm). It has been used in conjunction with other common dyes such as ATTO 488, ATTO 514, Alexa Fluor 488, 594, and 647 (Reitz et al., 2021).

We are interested in utilising a long-Stokes-shift fluorophore for in vivo *Drosophila* brain two-photon imaging alongside genetically encoded indicators, for example GCaMP, which fluoresce green with an emission peak wavelength of 510 nm (Nakai et al., 2001; Zhang et al., 2023) when excited by, for example, a 920 nm ultrashort pulse laser. Under single-photon conditions, ATTO 490LS has a similar excitation peak to GFP and its derived sensors like the GCaMPs, making ATTO 490LS a potential reference dye during neuronal imaging. However, ATTO 490LS’s two-photon properties have not yet been tested nor reported in literature. Here, we perform emission and excitation spectral analyses of streptavidin-conjugated ATTO 490LS to determine the dye’s suitability for two-photon imaging with systems that are configured with only a single fixed-wavelength 920 nm fibre laser, but have at least two detectors (Figure 1A).

**Figure 1.**
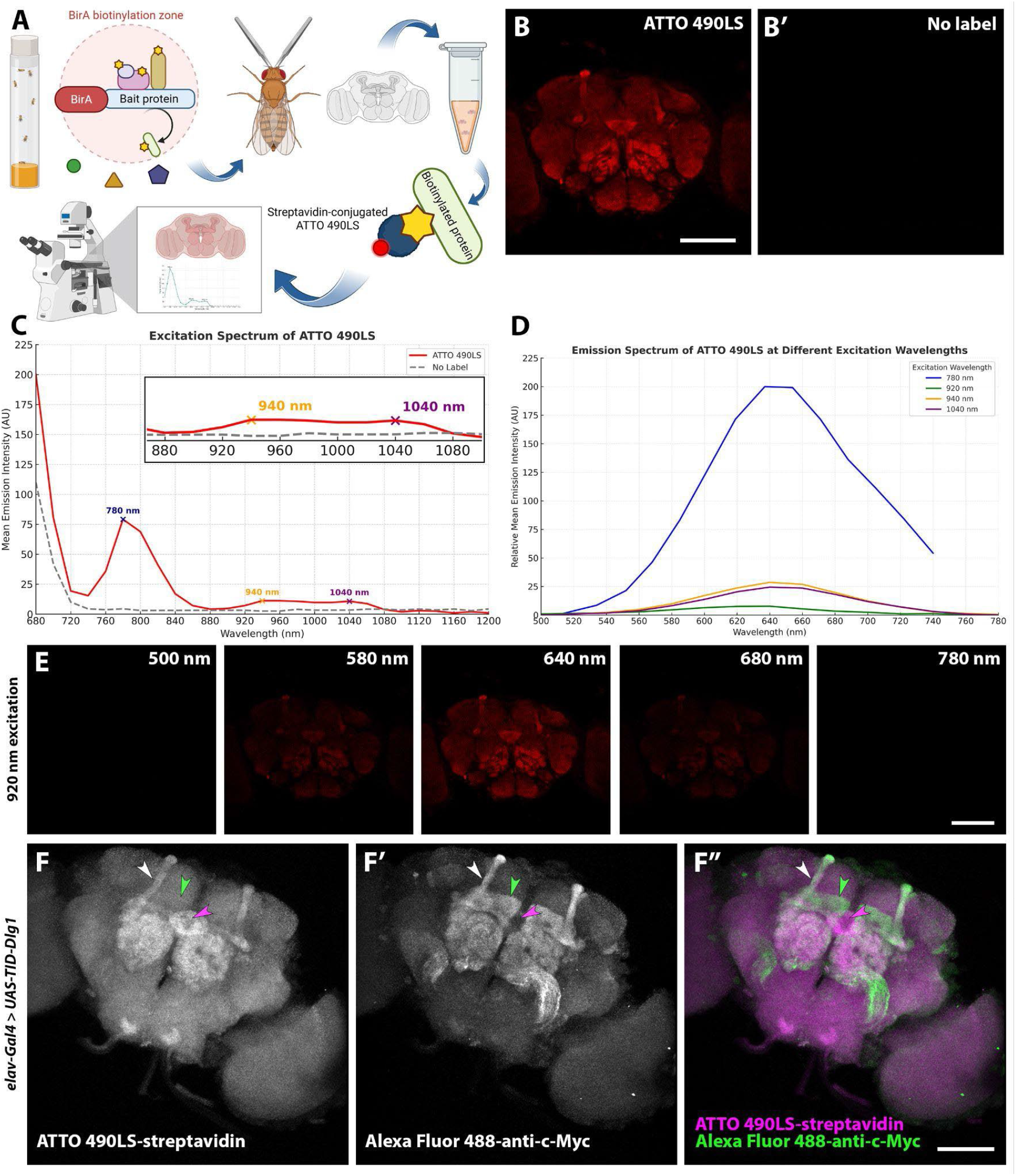
ATTO 490LS is excited by a 920 nm laser and is suitable for single-laser multicolour two-photon imaging. **(A)** Schematic diagram of the experimental workflow. **(B)** Representative image of *elav-Gal4>UAS-TID-Dlg1* fly brain, with ATTO 490LS staining at 780 nm excitation during the excitation scan. Scale bar = 100 μm. **(B′)** The identical experiment as **B**, but with no staining. **(C)** Normalised excitation spectrum of ATTO 490LS (red line) over 680 nm to 1200 nm compared with the emission intensity from an unlabelled brain (grey dashed line). Emission intensity from 680–740nm excitation is almost entirely attributed to autofluorescence. Emission intensity of ATTO 490LS peaks at 780 nm, 940 nm, and 1040 nm excitation, with normalised mean fluorescence intensities of 79.1, 11.1, and 10.71 (arbitrary units) respectively. Insert shows an enlarged view of the emission peaks at 940 nm and 1040 nm, which are 3–4-fold higher than the baseline emission intensity. **(D)** Normalised emission spectra of ATTO 490LS over 500 nm to 780 nm, at 780 nm (blue line), 920 nm (green line), 940 nm (orange line), and 1040 nm (burgundy line) excitation. Excitation at all four tested wavelengths gives emission peaks at 640 nm, with normalised mean fluorescence intensities of 200.0, 7.5, 28.1, and 24.1 (arbitrary units) respectively. **(E)** Respective images at various emission wavelengths of an ATTO 490LS-stained *elav-Gal4>UAS-TID-Dlg1* brain from the 920 nm excitation emission scan. Scale bar = 100 μm. **(F–F′′)** Maximum intensity projection images of an *elav-Gal4>UAS-TID-Dlg1* brain labelled with streptavidin-conjugated ATTO 490LS and anti-c-Myc tag antibody bound with Alexa Fluor 488 secondary antibody. Both fluorophores were simultaneously excited by the 920 nm laser at 3% power (30 mW). Detection of **(F)** ATTO 490LS-streptavidin signal with the red-filtered PMT at 1% gain, **(F′)** Alexa Fluor 488-anti-c-Myc signal with the green-filtered PMT at 5% gain, and **(F′′)** merged signals. Magenta arrowheads mark the ellipsoid body, an example structure labelled here only by ATTO 490LS. Green arrowheads mark the γ-lobe of the mushroom body, an example structure labelled here only by Alexa Fluor 488. White arrowheads mark the α-lobe of the mushroom body, an example structure double-labelled here by both ATTO 490LS and Alexa Fluor 488. Scale bar = 100 μm.

To determine the optimal two-photon wavelength for exciting ATTO 490LS, we first performed an excitation scan on stained fly brains, which involved measuring emission intensity while adjusting the excitation wavelength. The resulting emission intensity peaked at 780 nm, 940 nm, and 1040 nm excitation, with normalised relative mean fluorescence intensities of 79.1, 11.1, and 10.7 arbitrary units, respectively (Figure 1B, C). Emission was also detected in the 680–740 nm excitation range, however this could be almost entirely attributed to autofluorescence (Figure 1B′, C). The largest of these three emission peaks occurred at 780 nm excitation, thus indicating that is the most efficient laser wavelength within the tested range for maximum emission of the dye. As it is not necessary to excite at the global excitation maxima, and given that we are interested in assessing the dye for compatibility with GFP excitation, the second local maxima at 940 nm was promising as this is within close range of the commercially available fibre laser wavelength of 920 nm. Additionally, longer excitation wavelengths generate less autofluorescence signals, thus excitation at 920 nm is a more attractive option than at 780 nm.

To confirm that the wavelengths identified in the excitation scan are sufficient to excite ATTO 490LS and to verify the emission spectrum under two-photon illumination, we performed a series of emission scans. We measured the intensities of emission at different wavelengths while keeping the excitation wavelength fixed at one of the three local maxima. An emission scan was also done for 920 nm excitation. At all four tested excitation wavelengths, emission intensities peaked at 640 nm (Figure 1D, E), which is similar to the manufacturer’s reported emission wavelength of 661 nm under single-photon excitation. The normalised relative mean fluorescence intensities concurred with the excitation-scan readings, where 780 nm excitation led to the highest emission intensity (200.0 arbitrary units), while 920 nm excitation resulted in a lower emission intensity (7.5 arbitrary units). Taken together, results from both excitation and emission scans suggest that ATTO 490LS can be excited using a 920 nm two-photon laser and will emit at around 640 nm.

To assess the actual utility of ATTO 490LS with the type of microscope widely used for *Drosophila* two-photon imaging, we prepared ex vivo *elav-Gal4>UAS-TID-Dlg1* brains double-labelled with streptavidin-conjugated ATTO 490LS and anti-c-Myc tag primary antibody visualised with an Alexa Fluor 488 secondary antibody. Discs large 1 (Dlg1) is a crucial scaffolding protein, encoded by the *dlg1* gene in *Drosophila* (orthologous to the *DLG1*/*SAP97* gene in humans), and is a synaptic marker (Lahey et al., 1994; Woods et al., 1996). Here, *elav-Gal4* drives expression of the *UAS-TurboID-Dlg1* responder expressing the bait protein, which is tagged with the artificial peptide sequence Myc-tag, in post-mitotic neurons. Antibody staining against c-Myc will label TurboID-Dlg1, and is therefore expected to label generic neuronal scaffolding such as the post-synaptic density in the *Drosophila* brain. Meanwhile, staining with a streptavidin-conjugated dye will label any biotinylated proteins that are within a ∼15 nm proximity to TurboID-Dlg1 (Branon et al., 2018). The brains were imaged with an early model Thorlabs A-SCOPE two-photon microscope equipped with a fixed-wavelength 920 nm fibre laser. We observed strong ATTO 490LS signals through the red-filtered photomultiplier tube (PMT; bandpass filter at 570–640 nm) and were able to identify labelled structures such as the antennal lobes, ellipsoid body, and the α- and β-lobes of the mushroom bodies (Figure 1F).

To examine the simultaneous detection and distinguishability of red and green signals excited by a single 920 nm laser, we turned our focus to the signal collected by the green-filtered PMT (bandpass filter at 500–550 nm). For the most part, the anti-c-Myc and streptavidin signals coincided; for example, the α- and β-lobes of the mushroom bodies and antennal lobes were double-labelled by both dyes (Figure 1F–F′′, see white arrowheads; see also Extended Data). In some cases, the two fluorophores did not colocalise; for example, the ellipsoid body is exclusively labelled by ATTO 490LS (see magenta arrowhead), while the γ-lobes of the mushroom bodies are exclusively labelled by Alexa Fluor 488 (see green arrowheads). There is currently no explanation for such differences. Nevertheless, we establish that ATTO 490LS can be combined with Alexa Fluor 488 for duplexed imaging with a single 920 nm laser. Duplexed imaging with other fluorophores such as GCaMP should therefore also be possible.

In conclusion, we have characterised the two-photon optical properties of ATTO 490LS and demonstrated its utility for multicolour imaging with single-laser systems. The dye’s compatibility with standard 920 nm excitation substantially lowers technical barriers to duplexed two-photon microscopy, particularly for laboratories with legacy equipment. We aim to eventually engineer ATTO 490LS-conjugated ligands for HaloTag, SNAP-tag, and other tag proteins (Los et al., 2008; Marques et al., 2022) for in vivo chemogenetic labelling in transgenic *Drosophila* lines, enabling simultaneous imaging of GCaMP calcium transients (or other green sensors (Liu et al., 2022; Hao et al., 2024)) alongside a stable red dye for use as a structural marker or sensor denominator. Such a tool may have relative benefits compared to (or simply provide another option to) inherently fluorescent proteins with long Stokes shifts (Kim et al., 2022; Subach et al., 2022). By expanding the toolkit for multicolour two-photon imaging in *Drosophila* and other systems, this work creates new opportunities for integrating functional imaging with genetic and behavioural analyses, advancing our capacity to dissect neural circuit function in behaving animals.

## Methods

### *Drosophila* lines and rearing

A cross of female virgin ☿ pan-neuronal driver *elav-Gal4 (X)* (Bloomington *Drosophila* Stock Centre, Stock #458) and male ♂ TurboID direct fusion bait *UAS-TID-Dlg1 Drosophila melanogaster* flies were set up on standard food supplemented with dry yeast and flipped every 2–3 days onto fresh food with yeast. Offspring were transferred to food containing 50 µM biotin for 5 days, which led to biotinylation of proteins proximal to the TurboID-Dlg1 bait proteins (following the principles of BioID proximity labelling (Branon et al., 2018)), and allowed for subsequent binding of the streptavidin-conjugated dye to the biotinylated proteins (Figure 1A).

### Brain dissection and sample preparation

Biotin-fed *elav-Gal4>UAS-TID-Dlg1* flies were immobilised on ice and culled in ice-cold 70% ethanol for a few seconds. Flies were transferred to a SYLGARD 184-coated (Dow Corning) Petri dish, and brains were dissected in ice-cold 1× PBS using Dumont #5SF forceps (Fine Science Tools).

Dissected brains were fixed in 4% PFA in PBS-0.3% Triton X-100 for 20 minutes at room temperature and subsequently washed in PBS-0.3% Triton X-100 three times for 15 minutes at room temperature.

### ATTO 490LS staining

1 mg of desiccated streptavidin-conjugated ATTO 490LS (ATTO-TEC GmbH, product no. AD 490LS) was resuspended in 1 ml of MilliQ water to form a stock solution of 1 mg/ml. Stock solution was aliquoted and stored protected from light at -40°C.

Based on standard *Drosophila* brain immunohistochemistry protocols and a protocol for ATTO staining of 50 µm-thick fixed mouse brain slices (Reitz et al., 2021), samples were incubated in ATTO 490LS at a concentration of 1:200 in PBS-0.3% Triton X-100 for 48 hours in the dark at 4°C. The streptavidin-conjugated dye will bind to and label biotinylated proteins that are proximal to the TurboID-Dlg1 bait protein (Figure 1A). The dye was removed, and the samples were washed in PBS-0.3% Triton X-100 four times for 15 minutes at room temperature. Samples can be stored in PBS at 4°C for a few weeks before proceeding with mounting and imaging.

### Immunohistochemistry

For single-label antibody staining, an anti-c-Myc tag mouse monoclonal IgG_1_ primary antibody (Santa Cruz Biotechnology, catalogue no. sc-40) was added to the samples at a concentration of 1:200 in PBS-0.3% Triton X-100 for 48 hours at 4°C. For double-label staining, the primary antibody was added alongside ATTO 490LS. Samples were washed in PBS-0.3% Triton X-100 three times for 15 minutes at room temperature. Alexa Fluor 488 goat anti-mouse IgG secondary antibody (Invitrogen ThermoFisher, catalogue no. A11001) was added at a concentration of 1:500 and samples were incubated overnight in the dark at 4°C. Samples were washed in PBS-0.3% Triton X-100 four times for 15 minutes at room temperature. Samples can be stored in PBS at 4°C for a few weeks before proceeding with mounting and imaging.

### Emission and excitation spectral scanning

Samples were mounted on 1.0–1.2 mm-thick 25 × 75 mm glass slides (Biomedia) in PBS and sealed with 170 µm-thick 18 × 18 mm glass cover slips. Mounting media were excluded to prevent potential effects they may have on the excitation and emission scan readings. Excitation and emission scans were performed on a Leica TCS SP8 DIVE multiphoton microscope (Leica Microsystems, Wetzlar, Germany) equipped with a tuneable femtosecond pulsed multiphoton laser. Samples were imaged using an HC FLUOTAR L 25×/0.95 W Corr water immersion objective. Emission light was collected using Leica HyD photodetector.

For excitation scans (Λ, capital lambda), the laser wavelength was sequentially stepped from 680 nm to 1200 nm in 20 nm defined increments, while keeping the emission bandwidth between 500 nm and 685 nm. Power output delivered to the microscope remained constant at 65 mW ± 5 mW using the ‘Constant Power’ excitation control mode. For emission scans (λ, lowercase lambda), the excitation wavelengths were fixed at 780 nm, 920 nm, 940 nm, or 1040 nm. The emission detection bandwidth was sequentially stepped from 500 nm to 780 nm with a step size of 20 nm, with the exception of the scan at 780 nm excitation where the detection bandwidth was set to 500 nm to 740 nm (with a stepsize of 17–18 nm) due to limitations in collecting emission wavelengths that are too close to the excitation wavelength. Laser power setting (1%), detector gain (50), scan speed (200 Hz), and zoom (1×) were kept constant across all emission scans. Image acquisition and spectral detection were controlled using Leica LAS X software.

To quantify and compare emission intensities, each pixel of the subsequent 8-bit images was assigned a fluorescence intensity value from 0 to 255, where 0 represented no signal and 255 represented clipped or oversaturated signal (Shiffman, 2015), and a mean average of all pixels in the image were taken. Excitation and emission spectra graphs were generated using OpenAI by plotting normalised mean fluorescence intensity against excitation or emission wavelength, respectively.

### Two-photon imaging

Samples were mounted on 1.0–1.2 mm-thick 25 × 75 mm glass slides (Biomedia) in VECTASHIELD Antifade Mounting Medium (Vector Laboratories Inc.) and sealed with 170 µm-thick 18 × 18 mm glass cover slips. *Z*-stack image series were taken on two different Thorlabs microscopes (Thorlabs, NJ, USA).

The first setup was a factory-built Thorlabs multi-photon microscope (2010 model, referred to here as “A-SCOPE”) integrated with a Nikon Eclipse FN1 upright stand, a 7.9 kHz resonant/galvo scan head, a 1W FemtoFiber Ultra 920 laser (Toptica, Munich, Germany), a XLUMPLFLN 20× water immersion objective lens (Olympus, Tokyo, Japan), GaAsP PMT detectors, and ThorImageLS software (Thorlabs, NJ, USA). Laser power was set to 30 mW, unless stated otherwise. Emission light was collected by the green-filtered PMT, with a bandpass filter of 525/50 nm, and the red-filtered PMT, with a bandpass filter of 605/70 nm. Detector gains were set to 5% and 1% for the green- and red-filtered PMTs respectively, unless stated otherwise.

The second imaging setup was a Thorlabs B-SCOPE, equipped with CFI75 Apochromat 25×C W water immersion objective lens (Nikon, Tokyo, Japan) and Mai Tai eHP DS (Spectra Physics, CA, USA) running at 920 nm at 22.1 mW power. We used a set of 562LP and 635LP dichroic mirrors (Chroma) and 525/50 nm and 607/70 nm filters (Chroma) for the green and red channels respectively. Detector gains were set to 5% for the green and 1% for the red channel.

### Image processing

The two-photon image stacks were subjected to Gaussian blur 3D processing (X: 0, Y:0, Z:2) and maximum intensity projection processing in Fiji (Schindelin et al., 2012). For the purpose of visualising dual-channel images, signal collected by the green-filtered PMT was pseudo-coloured to green, while signal collected by the red-filtered PMT was pseudo-coloured to magenta. 3D rendered movies were created using the 3D Viewer plugin (Schmid et al., 2010) on Fiji.

The schematic diagram was created in BioRender. The figure was assembled in Adobe Photoshop.

## Reagents

| Animal Species | Genotype | Available From |
| --- | --- | --- |
| *Drosophila melanogaster* | *elav-Gal4 (X)* | Bloomington *Drosophila* Stock Centre (Stock #458) |
| *Drosophila melanogaster* | *UAS-TID-Dlg1* | Gift from Gunnar Poplawski and Keshika Ravichandran, manuscript in preparation. |

| Reagent/Antibody | Description | Available From |
| --- | --- | --- |
| ATTO 490LS | Streptavidin-conjugated | ATTO-TEC GmbH (product no. AD 490LS) |
| anti-c-Myc tag (9E10) | Mouse monoclonal IgG<sub>1</sub> | Santa Cruz Biotechnology (catalogue no. sc-40) |
| Alexa Fluor 488 | Goat anti-mouse IgG (H+L) | Invitrogen ThermoFisher (catalogue no. A11001) |

## Extended Data

**Extended Data Figure 1.**
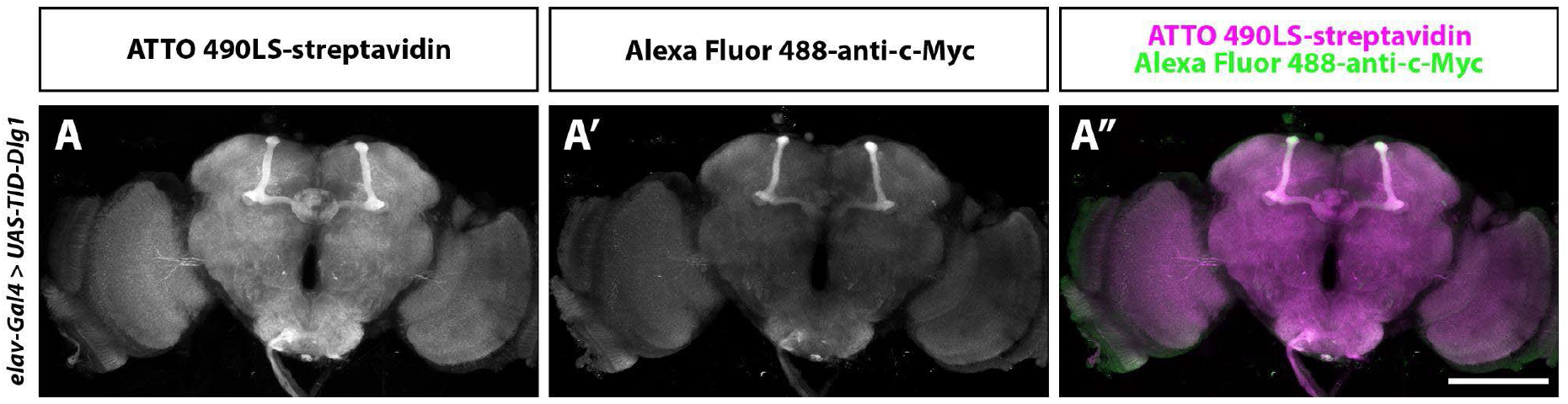
Duplexed two-photon imaging of *Drosophila* brain stained with ATTO 490LS and Alexa Fluor 488. Resource Type: Image **(A–A′′)** Maximum intensity projection images of an *elav-Gal4>UAS-TID-Dlg1* brain labelled with streptavidin-conjugated ATTO 490LS and anti-c-Myc tag antibody bound with Alexa Fluor 488 secondary antibody. Both fluorophores were simultaneously excited by the 920 nm laser at 22.1 mW power under the objective lens. *Z*-stack series was imaged from the posterior to the anterior side. Detection of **(A)** ATTO 490LS-streptavidin signal with the red-filtered PMT at 1% gain, **(A′)** Alexa Fluor 488-anti-c-Myc signal with the green-filtered PMT at 5% gain, and **(A′′)** merged signals. Scale bar = 100 μm.

**Extended Data Figure 2.**
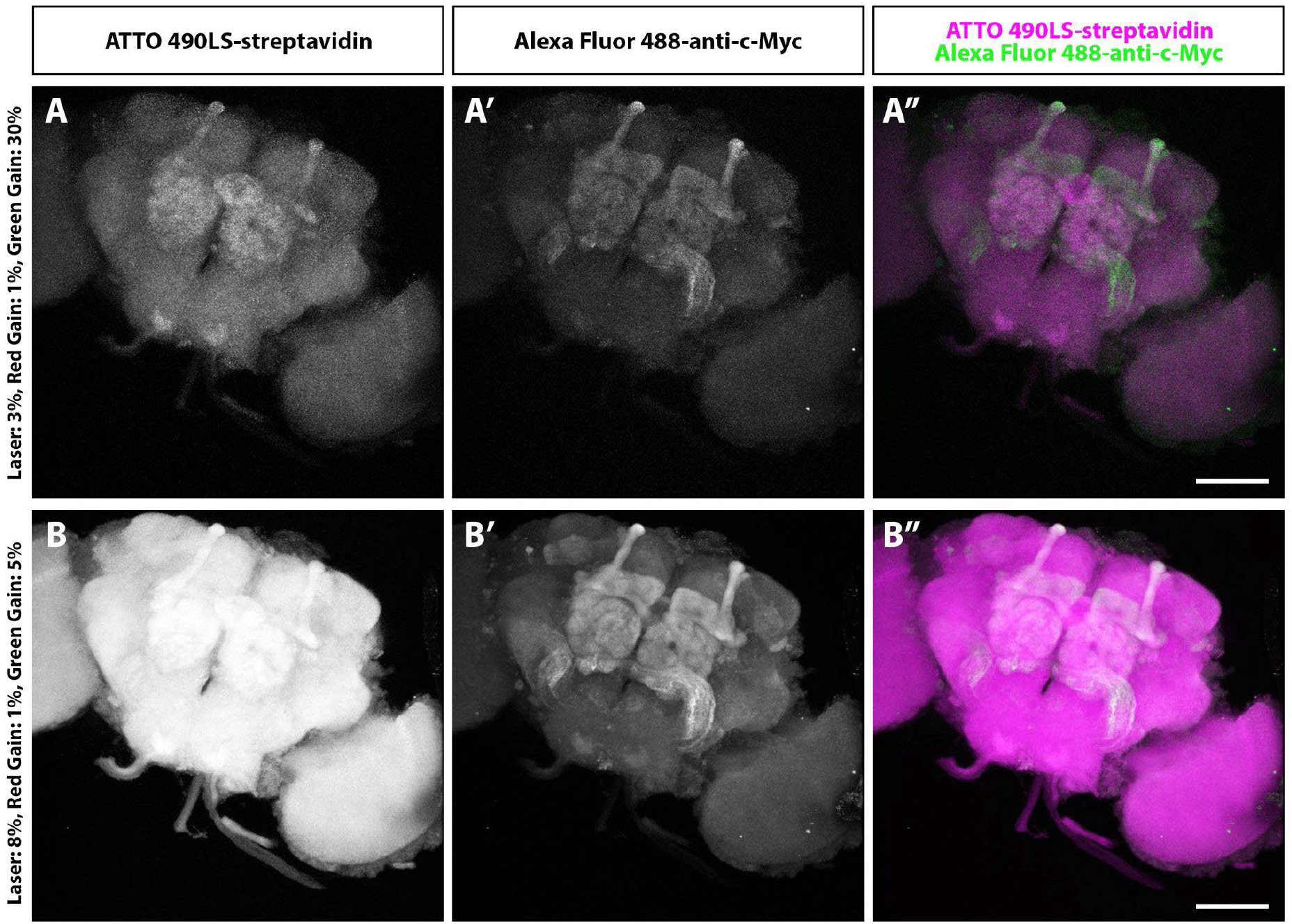
Duplexed two-photon imaging of *Drosophila* brain stained with ATTO 490LS and Alexa Fluor 488 at different laser powers and detector gains. Resource Type: Image **(A–A′′)** Maximum intensity projection images of an *elav-Gal4>UAS-TID-Dlg1* brain labelled with streptavidin-conjugated ATTO 490LS and anti-c-Myc tag antibody bound with Alexa Fluor 488 secondary antibody. Both fluorophores were simultaneously excited by the 920 nm laser at 3% power (30 mW). Detection of **(A)** ATTO 490LS-streptavidin signal with the red-filtered PMT at 1% gain, **(A′)** Alexa Fluor 488-anti-c-Myc signal with the green-filtered PMT at 30% gain, and **(A′′)** merged signals. Scale bar = 100 μm. **(B–B′′)** The same experiment as in **A–A′′**, but fluorophores were instead simultaneously excited by the 920 nm laser at 8% power (80 mW). Detection of **(B)** ATTO 490LS-streptavidin signal with the red-filtered PMT at 1% gain, **(B′)** Alexa Fluor 488-anti-c-Myc signal with the green-filtered PMT at 5% gain, and **(B′′)** merged signals. Scale bar = 100 μm.

**Extended Data Movie 1. *Z*-stack of two-photon image of *Drosophila* brain stained with ATTO 490LS and Alexa Fluor 488**.

Resource Type: Movie

*Z*-stack of Extended Data Figure 1.

**Extended Data Movie 2. 3D rendering of two-photon image of *Drosophila* brain stained with ATTO 490LS and Alexa Fluor 488**.

Resource Type: Movie

3D rendering of Extended Data Figure 1.

## Acknowledgements

We thank Gunnar Heiko Dirk Poplawski and Keshika Ravichandran (Programme in Neuroscience and Behavioural Disorders, Duke-NUS Medical School) for sharing the unpublished *UAS-TID-Dlg1* flies. We thank Sukanya Shyama Sundar (Multiphoton Core Facility — Multiphoton Microscopy Unit, Yong Loo Lin School of Medicine, National University of Singapore) for their technical support during the early stages of the project. We thank Stanislav Ott, Hyunsoo Shawn Je (Programme in Neuroscience and Behavioural Disorders, Duke-NUS Medical School), and Sangyu Xu (Institute of Molecular and Cell Biology, A*STAR) for insightful and productive discussions.

## Funding

K.Y.C., X.Z., D.D.S., and A.C-C. were supported by the OFYIRG20nov-0024 grant from the National Medical Research Council, Singapore, by grants MOE-T2EP30222-0018 and MOE-T2EP30223-0009 from the Ministry of Education, Singapore, and by a Duke-NUS Medical School grant. X.Z. and A.C-C. were supported by FY2022-MOET1-0001.

## Author Contributions

King Yee Cheung: conceptualisation, methodology, investigation, data curation, formal analysis, visualisation, writing — original draft, writing — reviewing and editing

Yue Wu: methodology, investigation, data curation, validation

Shu Ying Lee: conceptualisation, methodology

Xianyuan Zhang: investigation, writing — reviewing and editing

Masahiro Fukuda: investigation, writing — reviewing and editing

Danesha Devini Suresh: conceptualisation, writing — reviewing and editing

Adam Claridge-Chang: conceptualisation, supervision, funding acquisition, writing — reviewing and editing

## Notes

### Competing Interest Statement

The authors have declared no competing interest.

### Summary of Updates

Small wording edits in the main text, figure panel labels, and figure legend, in line with Reviewer/Editor comments.

