## Supplementary figures and images for "Two-photon characterisation of long-Stokes-shift dye ATTO 490LS for single-laser multicolour imaging"

### Extended Data Figure 1

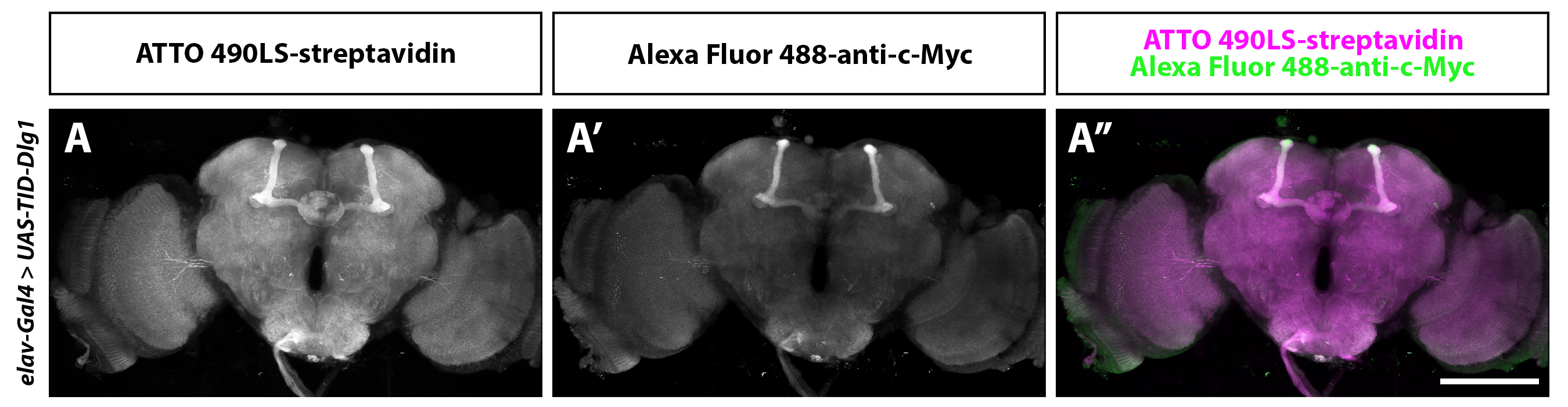

### Extended Data Figure 2

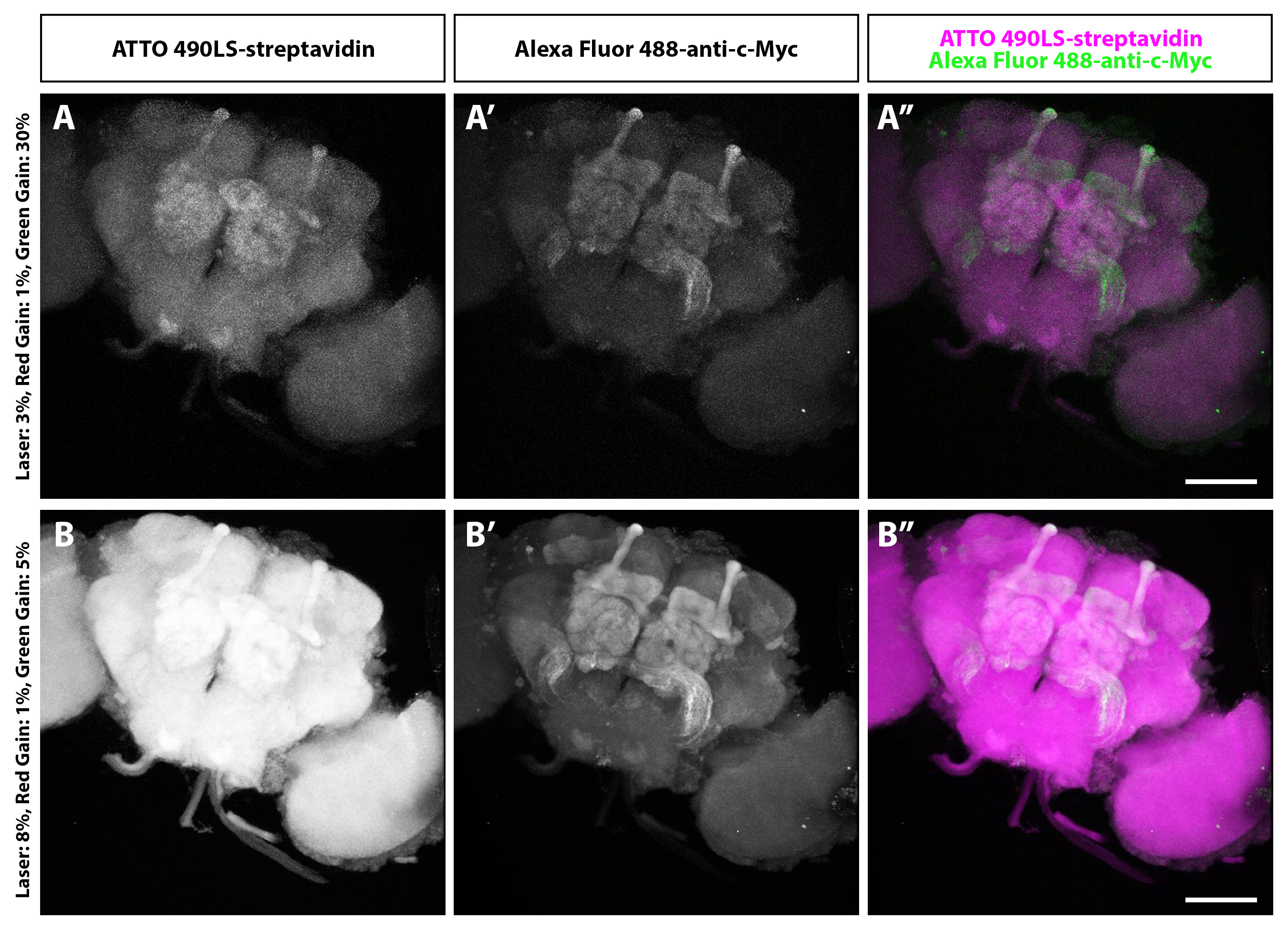
